# Severe inbreeding and gene loss in the historical and extant population of the critically endangered Devils Hole pupfish

**DOI:** 10.1101/2021.08.13.456274

**Authors:** David Tian, Bruce J. Turner, Christopher H. Martin

## Abstract

Small populations with limited geographic distributions are predicted to be threatened by inbreeding and lack of genetic diversity, both of which may negatively impact fitness and exacerbate population decline. One of the most extreme natural examples is the Devils Hole pupfish (*Cyprinodon diabolis*), an iconic and critically endangered species with the smallest known habitat range of any vertebrate. This imperiled species has experienced severe declines in population size over the last thirty years and suffered major, repeated bottlenecks in 2007 and 2013, when the population sunk to 38 and 35 individuals, respectively. Here we sequenced contemporary and historical genomes of Devils Hole and neighboring Death Valley and Ash Meadows desert pupfishes to examine the genomic consequences of small population size. We found extreme inbreeding (*F_ROH_* = 0.71 - 0.82) and increased genetic load in the Devils Hole pupfish. We also document unique fixed loss-of-function (LOF) alleles and deletions in genes associated with sperm motility, stress, and hypoxia within the extant Devils Hole pupfish population that likely reduce fitness. Comparisons between contemporary samples (2008 – 2012) and a genome sequenced from a 1980 formalin-fixed museum specimen suggest that inbreeding has increased 6% as the population has declined, but that many putatively deleterious variants have been segregating in the population since at least 1980. This includes a fixed early stop codon in *cfap43* (*n* = 8/8 samples), which is associated with sperm flagellum defects and causes infertility in humans and mice. Out of ninety-four unique deletions, fifteen were detected within 2 kb of annotated genes. Five have roles in physiological responses to hypoxia and mitochondrial activity, such as *redd1* (*n* = 7/7 samples), suggesting impaired hypoxia tolerance in this species despite the low oxygen concentrations of Devils Hole. We thus document one of the most extreme inbreeding events in a natural population and a set of candidate deleterious variants to inform management and potential genetic rescue in this conservation icon.

## Introduction

Due to the declining population sizes of many species and increasing isolation due to anthropogenic habitat fragmentation and climate change, understanding the extent and nature of genetic threats in small populations is essential for predicting and increasing population persistence and resiliency (Frankham 1995). Small and isolated populations are predicted to suffer from inbreeding depression, a reduction in fitness caused by the increased homozygosity of deleterious recessive alleles or overdominant loci (Charlesworth and Charlesworth 1999; Charlesworth and Willis 2009). Prolonged population decline can result in increased long-term extinction risk due to stochastic demographic events (Melbourne and Hastings 2008), reduced genetic variation for adaptation (Kohn *et al*. 2006), and decreased efficacy of purifying selection to overpower drift and purge deleterious variants (Kimura 1957) (reviewed by (Frankham 1995)).

Many examples of reduced genetic diversity and inbreeding depression have been documented in the wild such as Florida panthers (*Puma concolor*) (Johnson *et al*. 2010), Isle Royale wolves (*Canis lupus*) (Robinson *et al*. 2019), and mountain gorillas (*Gorilla beringei*) (Xue *et al*. 2015). Inbreeding associated population decline has been identified in the context of ancient bottlenecks (Palkopoulou *et al*. 2015; Rogers and Slatkin 2017) and on recent timescales (Robinson *et al*. 2019, 2021). Nevertheless, identifying inbreeding depression and its genetic basis in wild populations remains challenging due to small sample sizes and a lack of genomic resources. Recent studies of mountain gorillas (van der Valk *et al*. 2019) and Przewalski’s horses (Der Sarkissian *et al*. 2015) have also successfully leveraged museum specimens to measure temporal changes in inbreeding and genomic erosion. This is a powerful emerging approach because genomes from pre-decline museum specimens can serve as a baseline for estimating recent changes in inbreeding and genetic load (Bi *et al*. 2013; Díez-Del-Molino *et al*. 2018) and ultimately assist in restoring genetic diversity to a population (Bell *et al*. 2019).

Traditionally, conservation genetics has relied on neutral molecular markers to measure genetic variation in wild populations as a way of assessing population health and extinction risk, due to positive correlations between heterozygosity and fitness (Hansson and Westerberg 2002). Maintaining genome-wide genetic variation is crucial to preserving adaptive potential and preventing inbreeding depression in populations of conservation interest (Kardos *et al*. 2021).

However, several recent studies have suggested that genetic variation summary statistics do not necessarily accurately reflect population size or extinction risk and that we should also leverage genomics to assess functional genetic diversity and genetic load (Díez-Del-Molino *et al*. 2018; Teixeira and Huber 2021). Genomics also allows for more accurate measurements of individual inbreeding relative to pedigree approaches and the opportunity to infer the genetic basis of inbreeding depression (Kardos *et al*. 2016). Thus, a comprehensive understanding of the population and evolutionary dynamics of small population sizes in the wild from a genomic perspective is key to understanding the future of endangered populations and informing conservation management.

Here we leverage the unique evolutionary and demographic history of the Devils Hole pupfish (*C. diabolis*) to investigate how isolation and recent population decline have shaped inbreeding depression and genetic load. While many desert pupfishes are generally isolated with low population sizes, none are as extreme as *C. diabolis.* The habitat of *C. diabolis* is widely believed to be the smallest in the world for a vertebrate (3.5m x 22m) (Deacon *et al*. 1995) and the population has steadily declined since the late 1990s before reaching lows of 38 and 35 individuals in the spring of 2007 and 2013, respectively. Population viability analysis models in 2014 suggested that the median time to extinction is 26 years (Beissinger 2014). Although this species was previously believed to be isolated in Devils Hole for 10,000 – 20,000 years (Miller 1981), more recent genome-wide estimates suggest that Devils Hole may have been colonized most recently approximately 1-2 thousand years ago and suggest that gene flow among these desert oases is surprisingly common (Martin *et al*. 2016; Martin *et al*. 2017; Martin and Höhna 2018; Martin and Turner 2018). Furthermore, extremophile conditions include a nearly constant temperature of 34°C (Miller 1948; James 1969), no direct sunlight during the winter (Riggs and Deacon 2002) which limits primary production and nutrient availability (Wilson *et al*. 2001), and low dissolved oxygen levels near lethal limits for most fishes (2 – 3 ppm) (Gustafson, E.S. and Deacon, J.E. 1997; Riggs and Deacon 2002; Sackett *et al*. 2018). The continued persistence of *C. diabolis* in the hottest desert on earth in one of the most inhospitable habitats for fishes is nearly inconceivable.

Despite being one of the world’s most vulnerable species, genetic analyses have so far been limited to mitochondrial (Echelle and Dowling 1992), allozyme (Echelle and Echelle 1993), retrotransposon (Duvernell and Turner 1999), and RADseq (Martin *et al*. 2016; Martin and Höhna 2018) approaches. Here we used resequenced whole genomes of *C. diabolis* and several closely related *Cyprinodon* desert pupfishes to investigate how isolation and small population size influence inbreeding and genetic load on a genome-wide scale and across time in this iconic symbol of conservation.

## Results

### Geography and population structure

Principal components, admixture analyses, and genome-wide mean *F_st_* estimates (*C. diabolis*/*C. nevadensis* and *C. salinus* = 0.31) indicate the presence of population structure among Death Valley desert pupfishes, corroborating previous results (Martin *et al*. 2016) (Fig. 1). Differences in individual inbreeding both between and within species also provide evidence for population structure. The *C. diabolis* population is highly inbred while *C. nevadensis* and *C. salinus* populations generally exhibited low to intermediate levels of inbreeding, measured by the cumulative fraction of the genome made up of runs of homozygosity (ROHs) (Fig. 2A). ROHs made up less than 10% of the genome in our *C. nevadensis amargosae* and *C. nevadensis nevadensis* samples, both relatively undisturbed natural spring populations. *C. nevadensis shoshone* and *C. nevadensis pectoralis* tended to have a higher *F_ROH_* than other *C. nevadensis* species, consistent with previous surveys that called to attention their small population size and need for protection (Miller 1969) and the intensive management histories of these desert spring populations. Within a species, individuals from different spring populations often revealed contrasting levels of inbreeding, such as between the Indian Spring and School Spring populations of *C. nevadensis pectoralis*, indicating that isolation is strong between different springs with different management histories.

**Figure 1.**
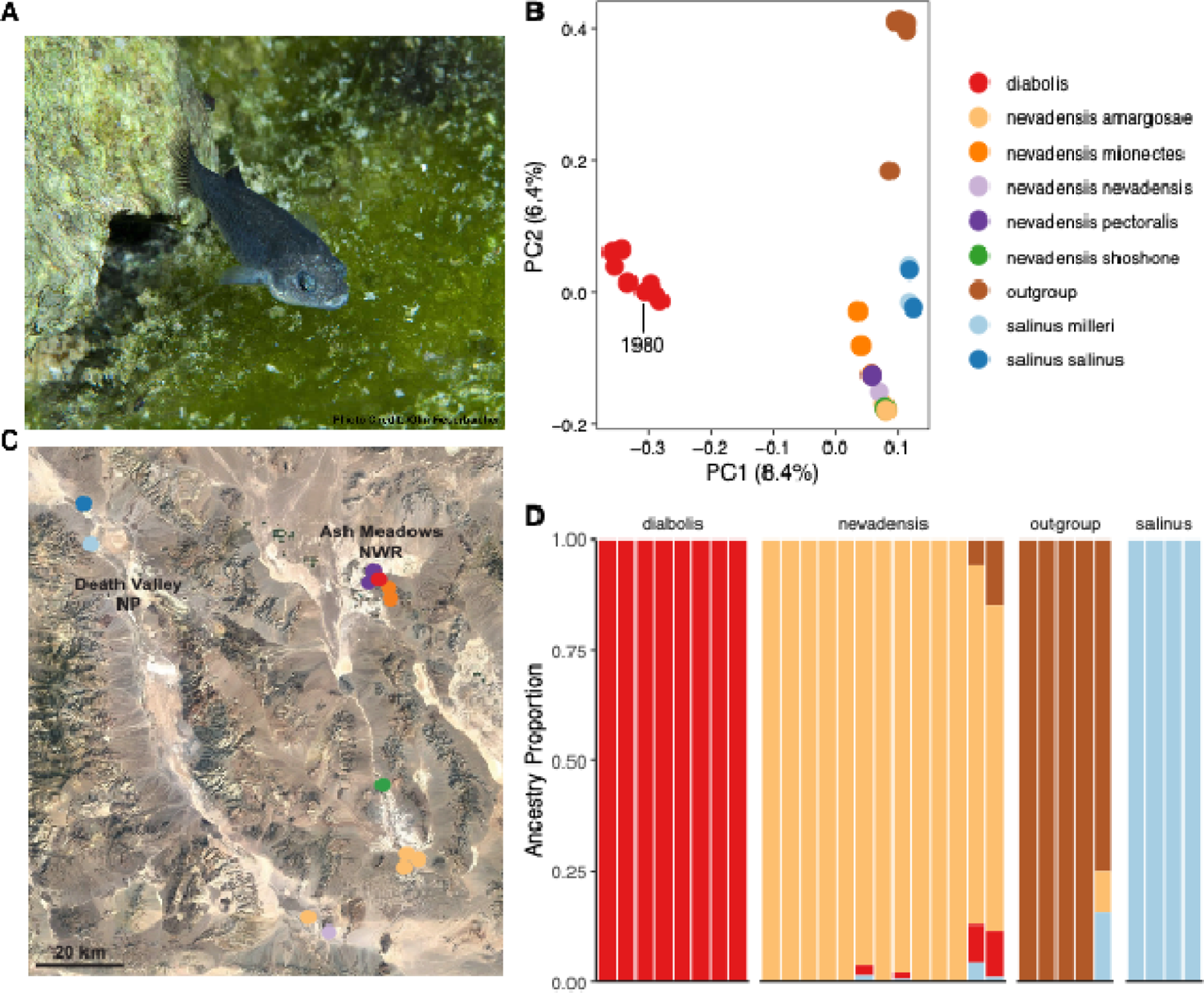
Geography and genetic structure of desert pupfishes A) Photo of male *C. diabolis* individual by Olin Feuerbacher. B). Principle component analysis of desert pupfishes revealing genetic population structure. C) Map of Death Valley NP and Ash Meadows NWR sampling locations. Colors in PCA designate individuals from different species. D) Ancestry proportions across individuals in Death Valley NP, Ash Meadows NWR, and outgroup desert pupfishes estimated from a LD-pruned SNP dataset in ADMIXTURE with k = 4.

**Figure 2.**
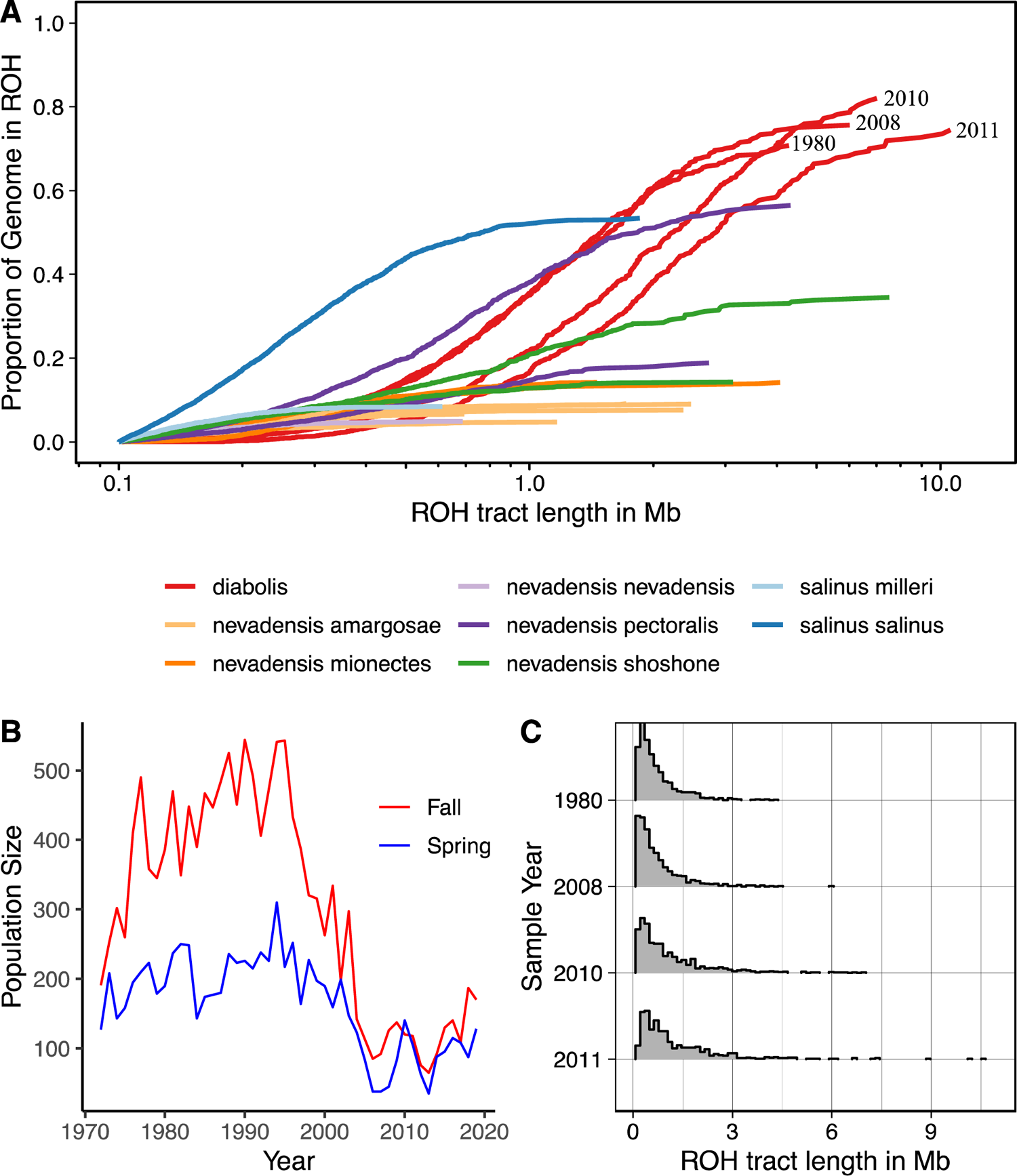
Extreme inbreeding in the Devils Hole pupfish A) *F_ROH_* measured as the cumulative fraction of the genome made up of runs of homozygosity (ROHs) at least 100 kb long. *F_ROH_* was calculated for all Death Valley pupfish individuals with at least 8X coverage. B) Biannual census counts of *C. diabolis* population size over time. Bottlenecks in 2007 and 2013 reduced the population size down to low as 38 and 35 individuals, respectively. C) Distribution of ROH lengths across *C. diabolis* samples from different time points. Greater values of *F_ROH_* among recent *C. diabolis* samples relative to 1980 can be primarily attributed to large ROHs (4 – 10 Mb) that are predicted to have arisen recently due to inbreeding within the last 1-3 generations, corresponding with the 2007 bottleneck.

### Severe inbreeding in Devils Hole pupfish

Strikingly, more than 70% of the *C. diabolis* genome consists of runs of homozygosity (ROHs), which is indicative of extreme inbreeding. Inbreeding can be identified and quantified through ROHs, which are long contiguous tracts of identical haplotypes inherited from a common ancestor (Kardos *et al*. 2016). For context, this level of inbreeding is far greater than other documented studies of notable, endangered species such as the Isle Royale wolves (Robinson *et al*. 2019) (< 50%), mountain gorillas (<40%) (van der Valk *et al*. 2019), and extant tigers (<50%) (Armstrong *et al*. 2021), with the exception of one population of lions from the Kathiawar Peninsula of India (90%) that had a population as low as 20 individuals in the early 20^th^ century (de Manuel *et al*. 2020).

The inbreeding observed within our 2008 - 2011 *C. diabolis* samples is ∼ 6% more severe than in our 1980 *C. diabolis* sample, which is consistent with the documented population decline that began in the mid 1990s and bottlenecks in 2007 and 2013 when the population size plummeted to 38 and 35, respectively (Beissinger 2014). While shorter ROHs indicate mating between distant relatives or longer-scale historical processes, long ROHs are indicative of recent inbreeding. We can estimate the mean number of generations back to the common ancestor of these homologous sequences based on the expectation that recombination breaks down long ROHs over time (Thompson 2013). We focused our attention on longer ROHs for which we have more confidence in timing and investigated long ROHs that are present in our 2008 – 2011 samples but not our 1980 sample. These long ROHs ranged from 4 – 10 Mb and are estimated to have originated only a few generations prior, suggesting that while the population began to decline a decade earlier, much of the additional inbreeding occurred more recently, due to the 2007 bottleneck. Our results illustrate that extreme isolation and prolonged small population size have driven *C. diabolis* to become one of the most inbred wild populations surviving today.

Recent population bottlenecks increased *F_ROH_*, but it was already extremely inbred prior to the recent population decline in the 1990s.

### Highest genetic load in Devils Hole pupfish

To measure genetic load, we used three summary statistics: the number of variants, derived alleles, and derived homozygous genotypes per individual, per site (Figure 3). We measured these statistics for synonymous, non-synonymous, and loss-of-function (LOF) mutations. *C. diabolis* had significantly more non-synonymous derived alleles (*P* = 0.0423) and derived homozygous genotypes than our *Cyprinodon* desert outgroups (*P* = 0.0258, Tukey’s HSD tests). Furthermore, *C. diabolis* had significantly more LOF variants than *Cyprinodon* desert outgroups (*P* = 0.0010, Tukey’s HSD test) and *Cyprinodon diabolis, nevadensis*, and *salinus* all had more LOF derived alleles than outgroups (*P* = 8.36 x 10^-5^; *P* = 0.0012; and *P* = 0.0229; Tukey’s HSD test, respectively). We found a similar pattern for LOF homozygous derived genotypes, although in this case *C. diabolis* also had significantly more LOF homozygous derived genotypes than *C. nevadensis* (*P* = 0.0036, Tukey’s HSD test).

**Figure 3.**
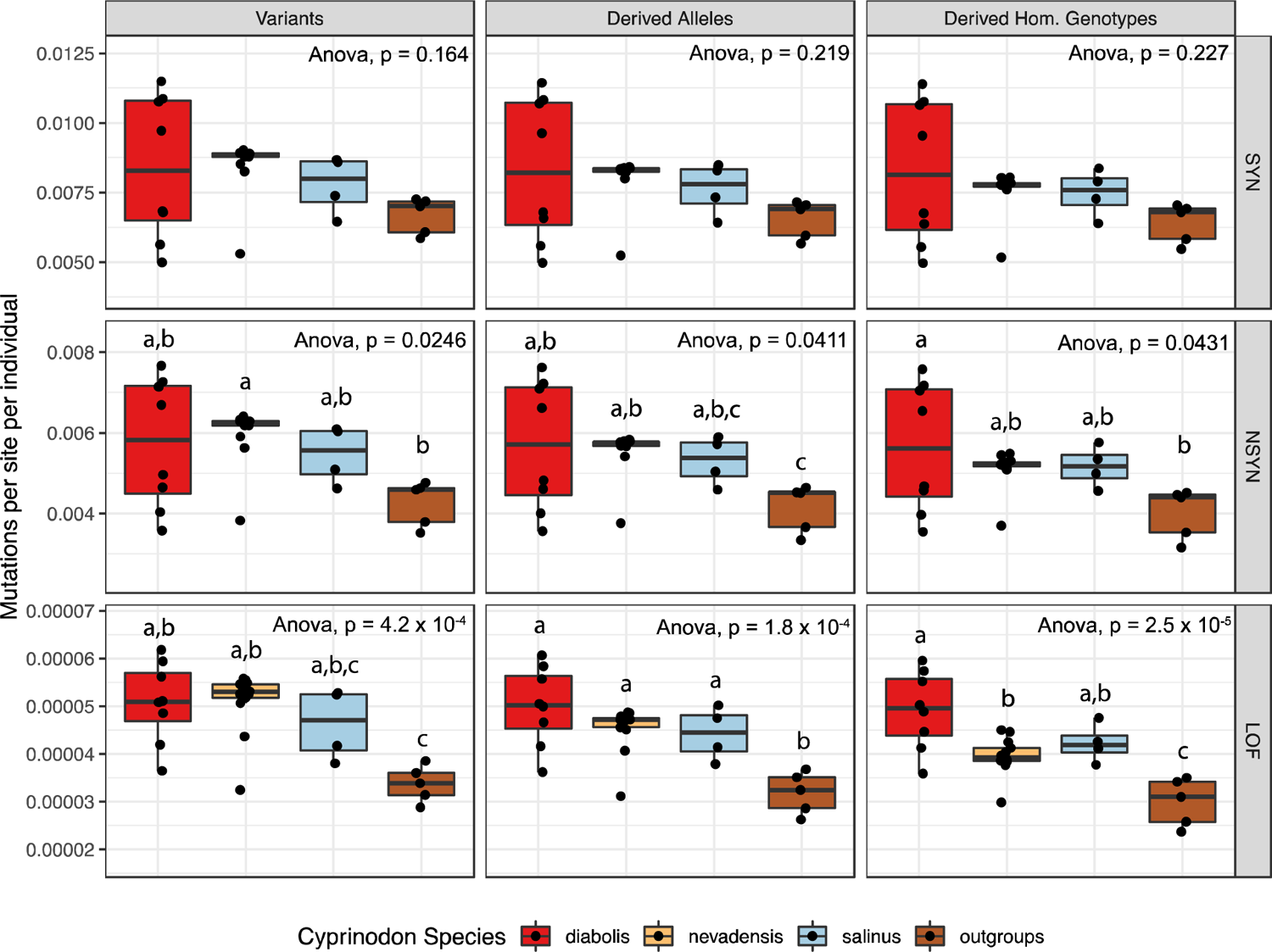
Genetic load across *Cyprinodon* species The number of synonymous, non-synonyomous, and loss-of-function mutations per site, per individual. Genetic load was measured in terms of three proxies: number of variants, number of derived alleles, and number of derived homozygous genotypes. The number of derived alleles per individual can be used to quantify load under an additive model while the number of derived homozygous genotypes can be used to quantify load under a recessive model. Results were scaled by the number of genotyped sites per individual to account for varying coverage. Loss-of-function mutations are those that encode a premature stop codon. Letters indicate which pairwise comparisons using a Tukey’s HSD test were significant.

### Fixed loss-of-function variants unique to Devils Hole

We searched for specific genetic variants that are predicted to be deleterious, may explain *C. diabolis* population decline, and can serve as a list of candidate deleterious variants to screen for potential genetic rescue (Table S1). We focused our attention on LOF SNPs and deletions unique to *C. diabolis*, which are expected to be deleterious and likely pose a threat by disrupting gene function.

**Table 1:** Unique deletions in *C. diabolis* tend to be involved in hypoxia Frequencies are based on samples for which there is enough coverage to determine whether the deletion is present or absent. Highlighted genes have functions related to hypoxia and mitochondrial stress. See extended data for details on intermediate scenarios where random spurious reads span the deletion or there is not enough coverage to make a definitive call of deletion presence or absence.

| Gene | Function | Location | Size<br>(bp) | Frequency |
| --- | --- | --- | --- | --- |
| *adamts15* | O-linked glycosylation | deletion in intron 7 | 55 | 5/5 |
| *apeh* | Destroys oxidatively-damaged proteins | exon 13 | 837 | 7/7 |
| *cnksr2* | MAPK/RAS signaling | intron | 53 | 7/7 |
| *cyp2a10* | Catalyzes the oxygenation of substrates | deletes exon 5/6 | 980 | 4/4 |
| *fbln7* | Angiogenesis in human umbilical vein<br>endothelial cells | 7 bp of exon 3, frameshift | 51 | 7/7 |
| *fzd1* | Regulates cumulus expansion genes,<br>required for normal female fertility in mice | 360 bp downstream | 66 | 6/6 |
| *nkx2_4* | Homeobox protein | deletes 3' UTR | 269 | 6/6 |
| *slc25a42* | Mitochondrial transporter of coenzyme A | start codon and TATA<br>box in promotor | 283 | 6/6 |
| *st6galnac1* | Glycosylation of proteins | deletes entire exon 4 and<br>large chunk of intron 3 | 2747 | 6/6 |
| *tmem206* | Acid-chloride channel, under selection in<br>Tibetans for adaptation to high altitude | intron 4 | 346 | 7/7 |
| *trim39* | Erythropoiesis in zebrafish | 254 bp of last exon<br>including stop codon, 3' | 854 | 5/5 |

**Table 1: Unique deletions in *C. diabolis* tend to be involved in hypoxia**
|  |  | UTR |  |  |
| --- | --- | --- | --- | --- |
| *redd1* | Suppresses protein translation to conserve energy | TATA box and hypoxia response element in promotor | 81 | 6/6 |
| *rnf220* | E3 ubiquitin ligase | 1313 bp downstream | 139 | 7/7 |
| *skil* | Component of SMAD pathway, which regulates cell growth and differentiation. | 1830 bp downstream | 230 | 5/5 |
| *slc8a1* | Required for normal embryonic heart development | 1477 bp downstream | 61 | 6/6 |

We found 16 predicted LOF SNPs that were primarily premature stop codons, including several within genes that potentially reduce fecundity and resistance to disease and stress (Table S1). These included *ora1*, a zebrafish olfactory receptor which is predicted to be a reproductive pheromone because it induces increased oviposition frequency (Behrens *et al*. 2014) and *trim25*, an ubiquitin ligase that promotes innate immune function in zebrafish (Jin *et al*. 2019). We also found a gained premature stop codon in *cfap43*, a protein involved in the conserved structure and function of the sperm flagellum axoneme and implicated in the multiple morphological abnormalities of the flagella (MMAF) infertility phenotype (Tang *et al*. 2017; Coutton *et al*. 2018). In nine men with MMAF that had homozygous mutations in *cfap43*, five had gained a premature stop codon (Coutton *et al*. 2018). Intriguingly, reduced sperm motility and abnormal sperm morphology is also an inbreeding phenotype found in Florida panthers (Johnson *et al*. 2010), East African and Indian lions (Wildt *et al*. 1987b), wild and captive bred cheetahs (Wildt *et al*. 1983, 1987a) and Dryas monkeys (van der Valk *et al*. 2020), which have a homozygous LOF mutation in *sept12* that causes a defective sperm annulus (Kuo *et al*. 2012).

We also searched for variants that may be responsible for recent increased inbreeding depression post 1980 by identifying variants that were heterozygous in our 1980 sample but were homozygous derived in our more recent samples after 2008 (Table S2). We focused our attention on LOF and non-synonymous variants. We found a premature stop codon in *dhx57*, a putative ATP dependent RNA helicase (UniProt 2008), and non-synonymous mutations in *arhgap18*, a GTPase-activating protein that modulates cell signaling (Maeda *et al*. 2011), and *galnt3*, a GalNAc transferase implicated in abnormal calcium phosphate deposits in various tissues (Sun *et al*. 2016). Moreover, we found two non-synonymous mutations that both occurred in *itprid2*, which encodes a protein that serves as a sperm surface antigen involved in cleavage of the fertilized oocyte (Javed and Naz 1992).

### Fixed deletions unique to Devils Hole

We searched for unique homozygous deletions in the *C. diabolis* genome that may negatively impact fitness. We identified 94 deletions unique to *C. diabolis* and focused our attention on the 15 deletions within 2 kb of any annotated gene. Surprisingly, 5 of the 15 deletions were involved in responses to hypoxia, mitochondrial activity, and oxidative stress (Fig. 4). Hypoxia leads to a wide suite of adaptive cellular responses that include angiogenesis and erythropoiesis, shifts to anaerobic metabolism, inhibition of translation, degradation of misfolded proteins, and regulation of nutrient use and cell fate (Lee *et al*. 2020).

We found an 81 bp deletion in the promoter of *redd1*, which is activated by HIF1 (DeYoung *et al*. 2008). This deletion knocks out the TATA box and a HRE recognition site where HIF1 is predicted to bind. During hypoxia, *redd1* inhibits mTORC1 (Brugarolas *et al*. 2004; DeYoung *et al*. 2008) through the TSC1/TSC2 tumor suppressor complex to conserve energy and prevent the accumulation of misfolded proteins. Intriguingly, upregulation of a homolog of REDD1 (*ddit4L*/*redd2*) has been implicated in adaptation to hypoxia in shortfin mollies (*Poecilia mexicana*) from H_2_S rich springs (Barts 2020) and high-altitude deer mice (*Peromyscus maniculatus*) (Rochette 2021 pers. comm.). Further evidence that *C. diabolis* is unable to properly inhibit protein translation through mTOR comes in the form of a fixed SNP that leads to an early stop codon in *tbc1d7*, a third core subunit of the TSC1/TSC2 complex whose knockdown leads to decreased association between TSC1 and TSC2 and thus increased mTORC1 signaling (Dibble *et al*. 2012).

Several deletions also likely result in either no protein transcribed or truncated non-functional proteins. *Apeh*, an enzyme acylpeptide hydrolase which destroys oxidatively damaged proteins, contains a fixed 837 bp deletion of exon 13 (Gogliettino *et al*. 2014; Riccio *et al*. 2015). *Trim39*, which in humans has been shown to regulate cell cycle progression and DNA damage responses, contains a fixed deletion of the last 254 bp of the last exon (Zhang *et al*. 2012b; a). However, *trim39* has undergone multiple rounds of duplications in several teleost lineages; in zebrafish it is known as *bloodthirsty* (*bty*) and is involved in erythropoiesis (Yergeau *et al*. 2005; van der Aa *et al*. 2009). There is also a 283 bp deletion containing the start codon for *slc25a42*, a mitochondrial transporter of coenzyme A (Fiermonte *et al*. 2009; Zallot *et al*. 2013). Mutations in *slc25a42* have been associated with mitochondrial myopathy and lactic acidosis in humans (Shamseldin *et al*. 2016; Iuso *et al*. 2019). *slc25a42* knockdowns in zebrafish cause severe muscle abnormalities that include dorsal curvature, bent tails, and early motor defects (Shamseldin *et al*. 2016).

## Discussion

We document extreme inbreeding in the critically endangered Devils Hole pupfish and identify a set of candidate deleterious structural and SNP variants to inform future genetic restoration. We show that *C. diabolis* is highly inbred, more so than all other Death Valley and Ash Meadows desert pupfishes and nearly all other known examples of inbreeding in the wild. By sequencing a historical museum specimen, we found that inbreeding has recently increased 6% from 1980 levels as the population has declined due to the presence of large (4 – 10 Mb) ROHs. Longer ROHs tend to have higher densities of deleterious alleles and larger fitness effects (Stoffel *et al*. 2021a). Although we were unable to directly measure fitness, the increase in inbreeding of 6% from 1980 to 2008 – 2012 likely resulted in a substantial reduction in fitness. Indeed, previous studies have suggested that increases in *F_ROH_* have strong negative effects on fitness; in Soay sheep an increase in *F_ROH_* by 10% resulted in a 60% decline in fitness (Stoffel *et al*. 2021b) while in helmeted honeyeaters, a 9% increase in homozygosity reduced lifetime reproductive success by 87-90% (Harrisson *et al*. 2019).

The already substantial amount of inbreeding seen as early as 1980 suggests that *C. diabolis* has likely frequently experienced population bottlenecks in the past prior to their recent population decline and bottlenecks. We know of at least one period of population decline that occurred in the late 1960s to early 1970s when increased groundwater extraction in Ash Meadows reduced water levels and jeopardized a shallow shelf in Devils Hole necessary for feeding and spawning (Deacon *et al*. 1995), leading to a landmark U.S. Supreme Court case over water rights (*Cappaert v. United States* 1976). While additional inbreeding has likely contributed to lower fitness and population decline, it is also important to note that population decline was amplified in 2004 by extreme rain that scoured the shallow spawning shelf in Devils Hole and resulted in the accidental trapping deaths of at least 72 individuals due to a flash flood (Beissinger 2014; Echelle and Echelle 2020). The lek-breeding system of *Cyprinodon* pupfishes also likely produces substantial reproductive skew every generation (Arnold 1972; Kodric-Brown 1977).

*C. diabolis* harbors higher genetic load than other desert pupfishes for nearly all classes of variation. However, load was generally not significantly different among *Cyprinodon diabolis / nevadensis / salinus* with the exception of the number of homozygous derived LOF genotypes, for which *C. diabolis* had significantly more than *C. nevadensis.* This implies that *C. diabolis* has a substantial genetic load especially if the LOF variants are recessive and strongly deleterious, and that extinction risk likely remains high (Kyriazis *et al*. 2021). The number of homozygous derived genotypes for LOF variants quantifies load under a recessive model (Henn *et al*. 2015, 2016), which is appropriate given that selection and dominance coefficients are generally inversely related (Simmons and Crow 1977; Agrawal and Whitlock 2012). Our evidence of higher genetic load in *C. diabolis* supports previous hypotheses of high load in this species due to its isolation and small population size, short generation time (Chernoff 1985; Deacon *et al*. 1995), and the observation that a refuge population of *C. diabolis* that was invaded by a few *C. nevadensis* individuals experienced rapid shifts in the gene pool to *C. nevadensis* alleles, leading to speculations of mutational meltdown (Lynch *et al*. 1995; Martin *et al*. 2012). Although recent studies have suggested that small populations generally have lower load (Xue *et al*. 2015; Robinson *et al*. 2018; Grossen *et al*. 2020), purging is expected to occur over long evolutionary timeframes and thus populations that have experienced recent severe population contraction are likely to have relatively high genetic load as deleterious variants reach fixation before purging removes them, particularly if prior population sizes were much larger (van der Valk *et al*. 2021).

### Degradation of hypoxia and reproductive pathways in C. diabolis

We found deleterious variants associated with reproduction and hypoxia and mitochondria stress genes. In particular, we found LOF and non-synonymous SNPs associated with sperm morphology and function (*cfap43, itprid2*). Deletions have thus far been rarely studied or quantified in conservation genetics. However, analysis of woolly mammoth genomes from different time points found that a sample dated close to the time of extinction had accumulated more homozygous deletions (Rogers and Slatkin 2017), suggesting that they may be an understudied genetic threat to endangered populations (Díez-Del-Molino *et al*. 2018). We found that unique, fixed deletions in *C. diabolis* tended to be associated with hypoxia and mitochondrial stress, a known environmental stressor in the system. Our results suggests that *C. diabolis* is poorly equipped to physiologically deal with the stress of low dissolved oxygen in Devils Hole and that putatively deleterious mutations in key genes likely hamper reproduction and population growth. Inbreeding depression often influences phenotypes such as survival, reproduction, and resistance to disease and environmental stress (Keller and Waller 2002); previous studies noted that *C. diabolis* has low fecundity (James 1969; Liu, R.K. 1969), low egg viability and juvenile survivorship (Deacon *et al*. 1995), and lays more eggs at lower temperatures (28 °C) compared to the higher constant temperature of 33 °C in Devils Hole (Burg *et al*. 2019). These results highlight the importance of investigating deletions and structural variants to more comprehensively understand deleterious variation in endangered populations.

### Adaptation to the Devils Hole environment?

However, we cannot rule out that some of these fixed variants are potentially the result of local adaptation, as it is extremely difficult to distinguish between selection and strong genetic drift in small populations because both processes leave similar signatures in allele frequencies (Leigh *et al*. 2021). One possibility is that repeated adaptations have accumulated in hypoxia pathways for the unique selective environment of Devils Hole relative to all other springs. However, we think it more likely that after the first LOF deletion disabled the entire pathway, additional LOF mutations accumulated without functional effects in a neutral manner. Future transcriptomic work will improve our understanding of how functional hypoxia pathways are in *C. diabolis*.

### Did mutational meltdown cause the recent population decline?

The DHP population began to decline in the 1990s from historical populations sizes of 200 – 500 individuals to population sizes of less than 100 each year for still unknown reasons. One possible explanation is that changes in environmental conditions, such as reduced nutrient inputs due to the elimination of wild sheep grazing around Devils Hole after the installation of razor wire fencing around the perimeter. Historical accounts of Devils Hole report thick mats of green macroalgae possibly due to ample manure, rather than the current ecosystem dominated by less-nutritious cyanobacterial mats and possibly decreased dissolved oxygen levels (Madinger *et al*. 2016). Alternatively, previous genetic analyses have speculated that the population has a high genetic load that has led to “mutational meltdown” based on dramatic shifts towards *C. nevadensis* ancestry in a *C. diabolis* x *C. nevadensis* hybrid refuge population (Martin *et al*. 2012). Our historical and modern analyses suggest that severe inbreeding existed before population decline in the 1990s, but that inbreeding and its negative impacts on the population’s fitness has likely been exacerbated due to recent population declines.

## Conclusion

Our study adds to a growing number of studies of inbreeding in wild populations in which deleterious mutations have been predicted from sequencing data (Robinson *et al*. 2019; Grossen *et al*. 2020). These studies also indicate that inbreeding depression is stronger in the wild than previously thought. The demographic history and genome-wide measures of genetic load and deleterious variation in the Devils Hole pupfish suggest that the population remains in danger. Our estimates of effective population sizes based on the harmonic mean of biannual census data (Ne^Spring^ = 122, Ne^Fall^ = 209) suggest that *C. diabolis* has an effective size far below suggested minimums for the maintenance of sufficient genetic variation for adaptive capacity (Franklin 1980; Reed *et al*. 2003; Traill *et al*. 2007). Further genomic sampling in the future will better inform our understanding of the mutation load and the genetic basis of inbreeding depression in this iconic conservation symbol.

## Methods

### Samples and sequencing

We sequenced 44 whole genomes that encompass *C. diabolis*, and many of the *C. nevadensis* and *C. salinus* Death Valley pupfishes spanning multiple independent springs, along with several further removed outgroups from various areas of California, Arizona, and Mexico. Of the 23 *C. diabolis* genomes, 8 were from historical museum samples, spanning 1937 – 1980 while the rest were collected between 2008 - 2012. Given the critically endangered status of *C. diabolis*, NPS and USFWS staff collected dead specimens found during routine checks in this period. *C. nevadensis, C. salinus,* and outgroup samples were collected in the early 1990s (Duvernell and Turner 1998). Samples were sequenced to 10x coverage on 150 PE runs using an Illumina Novaseq. Historical and several degraded *C. diabolis* tissues were prepared using Swiftprep library kits. For sampling dates, see extended data.

### Alignment and filtering

Raw reads were mapped from 44 individuals to the *Cyprinodon brontotheroides* reference genome (v 1.0; total sequence length = 1,162,855,435 bp; scaffold N50 = 32 Mbp) with bwa-mem (v 0.7.12; (Li 2013)). Duplicate reads were identified using MarkDuplicates and BAM indices were created using BuildBamIndex in the Picard software package ((http://picard.sourceforge.net; v 2.0.1). We followed the best practices guide recommended in the Genome Analysis Toolkit (v 3.5; (DePristo *et al*. 2011)) to call and refine our single nucleotide polymorphism (SNP) variant dataset using the program HaplotypeCaller. SNPs were filtered based on the recommended hard filter criteria (i.e. QD < 2.0; FS > 60; MQ<40; MQRankSum < - 12.5; ReadPosRankSum < −8 because we lacked high-quality known variants.

Variants in poorly mapped regions were removed with a mask file generated from the program SNPable ((Li 2009); k-mer length =50, and ‘stringency’=0.5). Variants were then filtered by minimum (3) and maximum (315) depth based on recommendations (Li 2014). Variants were additionally filtered to remove SNPs with a minor allele frequency below 0.05, genotype quality below 20, or containing more than 50% missing data across all individuals at the site using bcftools (v 1.6; (Li 2011)) and vcftools (v.0.1.15; (Danecek *et al*. 2011)). Our flexible threshold of 50% missing data was necessary given the degraded nature of several samples. Our final dataset contained 6,295,414 SNPs.

### Population structure

We pruned SNPs in strong linkage disequilibrium using the LD pruning function (--indep-pairwise 50 5 0.5) in PLINK (v 1.9; (Purcell *et al*. 2007)) leaving 1.6 million variants. We then characterized population structure with two approaches. First, we used PLINK to conduct principal component analysis. Second, we used ADMIXTURE (v.1.3.0; (Alexander *et al*. 2009)) to assign individuals to variable numbers of population clusters (K = 1-20). We used the subset parameter in PLINK to randomly select 100,000 SNPs for analysis. *F_st_* between *C. diabolis* / *C. nevadensis* and *C. salinus* was calculated genome-wide per variant site on the 6.3 million variant dataset using the weir-fst-pop function in vcftools (v.0.1.15; (Danecek *et al*. 2011)).

Several samples were excluded from downstream analyses due to a low percentage of reads mapping to the reference genome and properly paired reads, presumably due to degradation. These include 7 out of 8 historical samples sequenced with the exception of one high quality sample (DHP1980-5). Several contemporary samples (DHP54904, DHP54908, DHP54910, DHP54914) were also excluded due to low quality. DHP54909 was excluded because it did not cluster with the remaining nine *C. diabolis* samples after observing the PCA. DHP54908, DHP54909, and DHP54910 were all collected during the summer of 2009 which may have led to degradation due to the summer heat in Death Valley. The genotypes of unique fixed deleterious variants and deletions were typically invariant or missing in degraded samples. Missing deletions and invariant sites among these samples may be due to degraded samples being more susceptible to contamination by environmental pupfish DNA during the amplification step of sequencing. For DHP54920, a duplicate genome was sequenced, and we only included DHP54920_N2 (higher coverage) in these analyses. We removed DHP54911 due to too much missing data (> 89%).

Out of 8 historical museum samples only one sample from 1980 (DHP1980-5) was successfully sequenced. While reads were obtained from the remaining historical samples, these samples had a low percentage of reads mapping and properly paired reads and tended to be low coverage. This 1980 sample clusters with our high quality 2008 – 2012 samples.

### Population size

Census data for 1972 – Spring 2013 was acquired from Beissinger 2014. Data for subsequent counts was based on released NPS news reports. Variance effective population size was calculated as the harmonic mean of spring (Ne^Spring^) and fall (Ne^Fall^) population counts over time.

### Runs of homozygosity

Runs of homozygosity (ROHs) were identified using the BCFtools/ROH command, which uses a hidden Markov model to identify autozygous portions of the genome from genetic variation data (Narasimhan *et al*. 2016). For each species within the Death Valley species complex, we analyzed samples with at least 8X coverage. Samples were run individually and allele frequency was estimated from the AC and AN values within the INFO field of the VCF file. To calculate the cumulative fraction of the genome consisting of ROHs, the identified ROHs were sorted, summed, and divided by the total size of the genome (1,162,855,435 bp). ROHs with a size below 100 kb or a quality score below 50 were filtered out. We calculated the proportion of the genome that was made up of ROHs at least 100 kb long (*F_ROH_*) for Death Valley samples with at least 8X coverage.

### Timing of inbreeding

We estimated the mean number of generations back to the common ancestor of homologous sequences with the following equation: g = 100 / 2*ROH length (cM), where g is the number of generations (Thompson 2013). Here we assume an average recombination rate across the genome of ∼ 4.6 cM/Mb based on an estimated genetic map length of 5330 cM (St. John *et al*. 2021) and genome length of 1.16 Gb in *C. brontotheroides* (Richards *et al*. 2021).

### Measuring genetic load

We categorized mutations in coding regions with regards to their effect on the amino acid sequence and whether the alleles were ancestral or derived with respect to the *C. brontotheroides* genome as the reference outgroup. We used Snpeff (Cingolani *et al*. 2012) to identify variants as either loss-of-function (LOF) or non-synonymous (NSYN) to measure genetic load. LOF variants were conservatively defined as SNPs that resulted in a premature stop codon following (Robinson *et al*. 2016, 2018), which are expected to be less prone to misannotation (de Valles-Ibáñez *et al*. 2016). We did not consider splice variants due to possible transcript differences across *Cyprinodon* species. We also did not include stop loss nor start loss variants in our measurements of load as stop loss variants are expected to have less important phenotypic effects (Richards *et al*. 2015) and start loss variants are difficult to assess due to potential alternative translational initiation at other start sites. Genetic load was measured in terms of three proxies: number of variants, number of derived alleles, and number of derived homozygous genotypes.

The number of derived alleles per individual can be used to quantify load under an additive model while the number of derived homozygous genotypes can be used to quantify load under a recessive model. Results were scaled by the number of genotyped sites per individual to account for varying coverage. We assessed whether differences in measures of genetic load among the four species groups were significant using an ANOVA in R and used pairwise Tukey’s HSD tests in R to test whether inter-species measurements of load differed.

### Unique deleterious variants

We used the following criteria to identify unique putatively deleterious variants: (i) homozygous, (ii) predicted to be loss-of-function or deleterious based on a Snpeff predicted impact of “HIGH” (Cingolani *et al*. 2012) (start codon loss, stop codon gain, stop codon loss, splice acceptor site variant, splice donor site variant) and (iii) present in at least our five higher quality *C. diabolis* samples (including our 1980 sample) and absent in all other non-*C. diabolis* samples in our dataset. To find unique variants that may be responsible for increased inbreeding depression post 1980, we also identified loss-of-function and non-synonymous variants that were invariant in *C. nevadensis*, *C. salinus*, and outgroups, but heterozygous in DHP1980-5 and homozygous derived in at least the higher quality more recent 2008-2012 *C. diabolis* samples.

### Unique deletions

We identified structural variation in the form of deletions that were unique to *C. diabolis* using DELLY (v0.8.3) (Rausch *et al*. 2012) following similar criteria as above (i and iii). We searched for deletions that were present in DHP1980-5, DHP54903, DHP54913, DHP54917, or DIAB54919, and not present in non-*C. diabolis* samples. Deletions that were exceptionally large and presumably artifacts were checked for accuracy in IGV and subsequently removed. We focused our attention on deletions within 2 kb of an annotated gene, based on annotations of the *C. brontotheroides* genome (Richards *et al*. 2021). Deletions were confirmed to be unique to *C. diabolis* by aligning bam files spanning the deletion in both *C. diabolis* and non-*C. diabolis* samples and confirming presence or absence, respectively. We analyzed whether deletions spanned exons, introns, or regulatory regions by blasting deleted sequences against the C_variegatus-1.0 (GCF_000732505.1) assembly on Ensembl (release 102; (Howe *et al*. 2021)).

## Author Contributions

DT wrote the manuscript and conducted all analyses. BJT and CHM collected the samples. DT and CHM contributed to the development of ideas presented in the study and revised the manuscript. All authors edited and approved the manuscript.

## Data and materials availability

Genomic sequence data will be archived at NCBI. Additional data will be deposited on the Dryad Digital Repository. All scripts are available on Github (https://github.com/tiandavid/).

## Acknowledgements

We thank members of the Martin lab for valuable comments and discussions; Olin Feuerbacher for providing a photo of *C. diabolis*; T. Echelle for providing specimens of *C. n. pectoralis* from Indian Spring; Lee Simons, U.S. Fish and Wildlife Service, and National Park Service for *C. diabolis* samples and permits; and the University of California, Berkeley for computational resources. This work was funded by the U.S. Fish and Wildlife Service, National Park Service, National Science Foundation DEB CAREER grant #1749764, National Institutes of Health grant 5R01DE027052-02, and the University of California, Berkeley to CHM. DT was supported by a National Science Foundation Graduate Research Fellowship (DGE 1752814).

## Supplemental Figures and Tables

**Table S1.** Unique, loss-of-function and deleterious variants in C. diabolis Loss-of-function variants that are invariant (0/0 or ./.) in *C. nevadensis*, *C. salinus*, and outgroups, but variant in at least higher quality *C. diabolis* samples. The frequencies of variants among *C. diabolis* individuals with genotype information at the given site are shown. DHX57 is heterozygous in our 1980 sample. See extended data for more information on the specific genotypes of individuals.

| Scaffold | Position | Ref/SNP | Variant Class | Gene | *C. diabolis*<br>Frequency |
| --- | --- | --- | --- | --- | --- |
| 1 | 25026331 | G/T | Gained stop codon | *cfap43* | 8/8 |
| 3 | 5977284 | G/A | Gained stop codon | *lamb1* | 6/6 |
| 4 | 13853843 | C/A | Gained stop codon | *herc3* | 6/6 |
| 20 | 7768 | G/A | Splice acceptor site | *ankle2* | 6/6 |
| 21 | 38986750 | G/A | Gained stop codon | *git1* | 7/7 |
| 26 | 9468645 | G/A | Gained stop codon | *dhx57* | 8/8 |
| 27 | 23697776 | T/A | Gained stop codon | *acsf2* | 6/6 |
| 34 | 34643729 | G/A | Lost start codon | *ora1* | 6/6 |
| 43 | 10030715 | G/A | Gained stop codon | *rims1* | 5/5 |
| 43 | 30333450 | G/A | Gained stop codon | *protein of unknown function* | 6/6 |
| 47 | 9410043 | G/T | Gained stop codon | *neoverrucotoxin subunit beta* | 6/6 |
| 53 | 23074581 | A/C | Lost stop codon | *cyth3* | 7/7 |
| 55 | 6497450 | T/A | Lost start codon | *acan* | 5/5 |
| 597 | 2816 | A/T | Gained stop codon | *tbc1d7* | 6/6 |
| 7768 | 3929 | C/T | Splice donor site | *b4galnt1* | 6/6 |
| 10811 | 499 | G/A | Gained stop codon | *trim25* | 5/5 |

**Table S2.** **Unique, putative variants associated with inbreeding depression in C. diabolis** Loss-of-function and non-synonymous variants that are invariant (0/0 or ./.) in *C. nevadensis*, *C. salinus*, and outgroups, but heterozygous (0/1) in DHP1980-5 and homozygous derived (1/1) in at least the higher quality more recent 2008-2012 *C. diabolis* samples. The frequencies of variants among *C. diabolis* individuals with genotype information at the given site are shown. Note that there are two separate non-synonymous variants present in *itprid2*. See extended data for more information on the specific genotypes of individuals.

| Scaffold | Position | Ref/SNP | Variant Class | Gene | Contemporary *C. diabolis* Frequency |
| --- | --- | --- | --- | --- | --- |
| 7 | 20355383 | G/C | Non-synonymous | *arhgap18* | 4/4 |
| 26 | 9468645 | G/A | Gained stop codon | *dhx57* | 7/7 |
| 37 | 1901551 | G/A | Non-synonymous | *Protein of unknown function* | 4/5 |
| 52 | 1780129,<br>1780297 | C/T, G/A | Non-synonymous | *itprid2* | 6/6, 6/6 |
| 52 | 12629951 | G/A | Non-synonymous | *galnt3* | 4/4 |

